# Deep mutational scanning to predict antibody escape in SARS-CoV-2 Omicron subvariants

**DOI:** 10.1101/2022.12.02.518937

**Authors:** Mellissa C Alcantara, Yusuke Higuchi, Yuhei Kirita, Satoaki Matoba, Atsushi Hoshino

## Abstract

The major concern of COVID-19 therapeutic monoclonal antibodies is the loss of efficacy to continuously emerging SARS-CoV-2 variants. To predict the antibodies efficacy to the future Omicron subvariants, we conducted deep mutational scanning (DMS) encompassing all single mutations in the receptor binding domain of BA.2 strain. In case of bebtelovimab that preserves neutralization activity against BA.2 and BA.5, broad range of amino acid substitutions at K444, V445 and G446 and some substitutions at P499 and T500 were indicated to achieve the antibody escape. Among currently increasing subvariants, BA2.75 carrying G446S partly and XBB with V445P and BQ.1 with K444T completely evade the neutralization of bebtelovimab, consistent with the DMS results. DMS can comprehensively characterize the antibody escape for efficient and effective management of future variants.

## Introduction

The Omicron variant demonstrated the tremendous escape from neutralization of vaccinated and convalescent serum samples and clinically used antibody drugs(1–3). Omicron subvariants keep obtaining mutations in the receptor binding domain (RBD) and the resultant accumulation of mutations induced the resistance to the majority of therapeutic antibodies. As of September, 2022, only bebtelovimab remains efficacy against major subvariants, BA.2, and BA.5 among clinically-used therapeutic antibodies(4–6).

To predict the neutralization activity against future variants, we performed deep mutational scanning (DMS) with the library encompassing all single amino acid substitutions in the BA.2 RBD. The library was expressed in the context of full-length spike on the cell surface and antibody neutralization was reproduced in the inverted infection assay where ACE-harboring viruses infected spike-expressing cells(1). This DMS identified K444, V445, and G446 as the critical sites for effective neutralization of bebtelovimab. Consistent with this result, currently increasing subvariants, XBB and BQ.1(7) are resistant to bebtelovimab due to V445P and K444T mutations, respectively.

## Results

### Deep mutational scanning for escape from bebtelovimab in BA.2

We performed deep mutational scanning (DMS) to determine how all single amino acid substitutions impact the susceptibility of BA.2 to bebtelovimab, the clinically used drug remaining neutralization activity against BA.2 and BA.5(4–6). The mutant library encompassed all 20 single amino acid substitutions in the RBD (F329 to C538) of BA.2 spike (Figure 1a). The plasmid library was expressed on human Expi293F cells in a manner where cells typically acquire no more than a single variant(1). The cells were infected with ACE2-harboring green fluorescent protein (GFP) reporter virus that was pre-incubated with bebtelovimab. The mutant spike escaping from the antibody allowed the infection of ACE2-harboring virus and the cells obtain GFP fluorescence, whereas mutations with preserved susceptibility to the antibody or compromised infectivity prevented the infection. Both infected GFP-positive cells and control GFP-negative cells were harvested with fluorescence-activated cell sorting (FACS). The RNA was extracted and underwent deep sequencing (Figure 1b). The DMS library was divided into 3 units and pooled after plasmid construction. Sequence samples were prepared with primers specific to each unit and mutations were identified by direct sequencing of the whole unit. The escape value was defined as the ratio of GFP positive read-count to the negative read-count and normalized by the value of wild-type(1). The spike protein–expressing cells constituted 20-30% of transfected cells, which was suitable to avoid multiple library induction(8). ACE2-harboring virus infected ~15% of transfected cells and bebtelovimab was titrated to reduce the infection rate to ~3% (1) (Figure 1c). DMS experiments were performed in duplicate and exhibited good reproducibility, R^2^ (coefficient of determination) = ~0.4796, as was the case in previous our and other studies (Figure 1d)(1, 9, 10).

**Figure 1.**
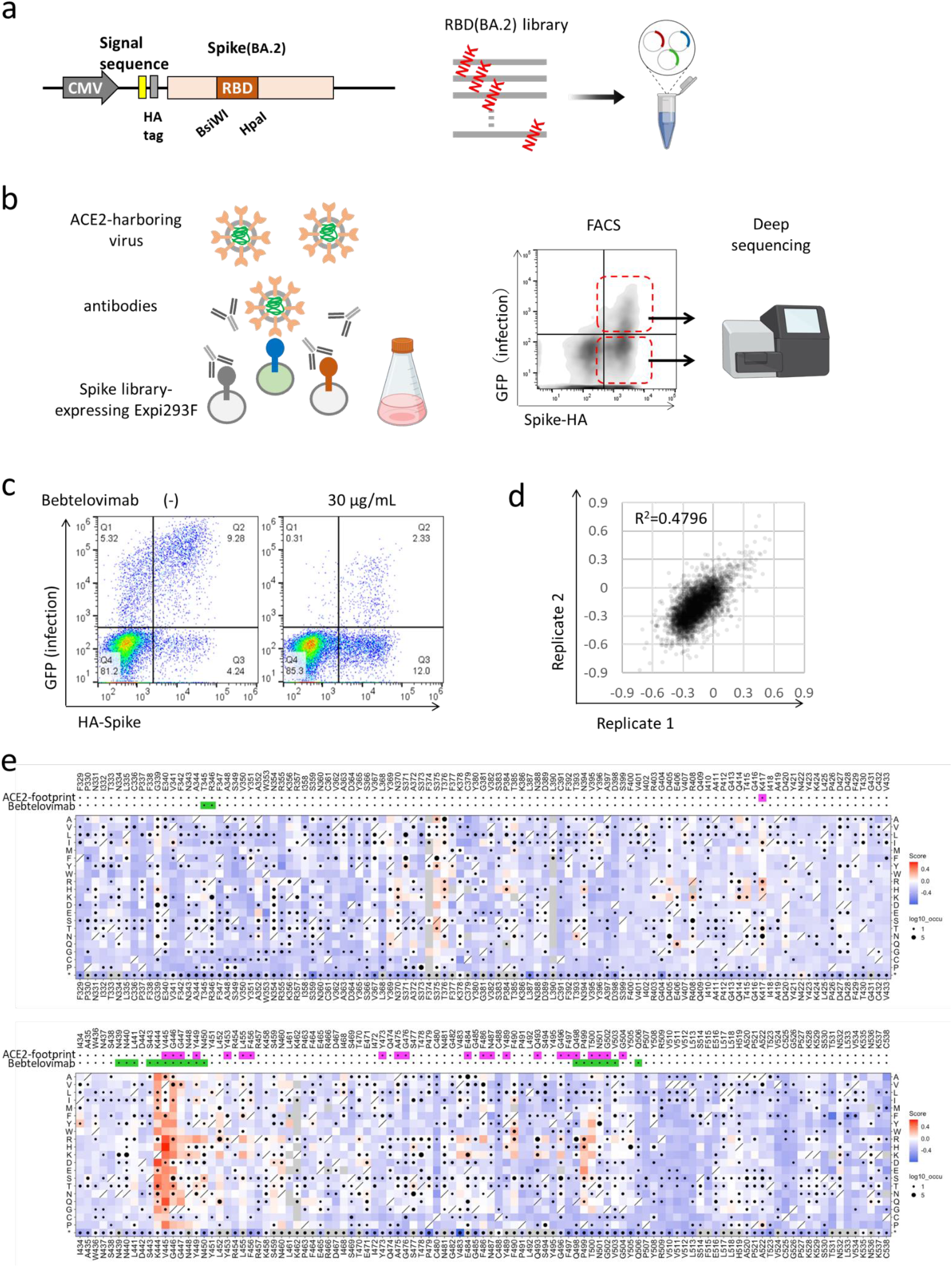
Deep mutational scanning identified single-residue mutation to induce escape from bebtelovimab. (**a**) A design of the deep mutational scanning (DMS) library for the receptor binding domain of BA.2 strain. (**b**) A schematic of the DMS approach to evaluate escape from neutralizing antibodies in the inverted infection assay. FACS, fluorescence activated cell sorting. (**c**) The experimental setting of library transfection and psuedovirus infection rate. (**d**) The correlation in mutation effects on the alteration of escape from bebtelovimab across replicates. (**e**) The heatmap illustrating how all single mutations affect the escape from bebtelovimab. Squares are colored by mutational effect according to scale bars on the right, with blue indicating deleterious mutations. Squares with a diagonal line through them indicate BA.2 strain amino acid. Black dot size reflects the frequency in the virus genome sequence according to GISAID database as 17^th^ Sep 2022. ACE2 and bebtelovimab footprints are highlighted in magenta and light green, respectively. Footprints on RBDs were defined according to 5A ° distance from ACE2- or monoclonal antibody-contacting residues(14).

The BA.2 based DMS indicated that broad range of amino acid substitutions at K444, V445 and G446 and some substitutions at P499 and T500 contribute to the escape from bebtelovimab, consistent with previous reports(11, 12) (Figure 1e). These sites were fairly unchanged before the emergence of the Omicron. However, Omicron subvariants obtained further wide variety of mutations including substitutions at K444, V445 and G446.

### XBB and BQ.1 evade bebtelovimab neutralization

We next evaluated the neutralization activity of bebtelovimab against major Omicron subvariants (Figure 2a). In the neutralization assay with pseudotyped lentivirus, BA.2, BA.2.12.1, BA.4/5, BA.4.6 are susceptible to bebtelovimab, whereas, BA.2.75 modestly and XBB and BQ.1 dramatically obtains resistance (Figure 2b). Another study also reported that bebtelovimab had lower neutralization activity against BA.2.75 albeit preserved activity against BA.1, BA.2, and BA.4/5(13). BA.2.75, BA.2-derived subvariant, additionally obtained D339H, G446S, and N460K in the RBD and the result of DMS indicated that G446S is responsible for reduced neutralization sensitivity against bebtelovimab. XBB and BQ.1 carry V445P and K444T, respectively, and these mutations are indicated to obtain the strong escape in the DMS. Although K444, V455, and G446 are the responsible region for bebtelovimab neutralization, not all amino acid substitutions provide the escapability. Especially, V445I, V445L and G446D are observed among circulating SARS-CoV-2 isolates and maintain sensitivity against bebtelovimab (Figure 1e).

**Figure 2.**
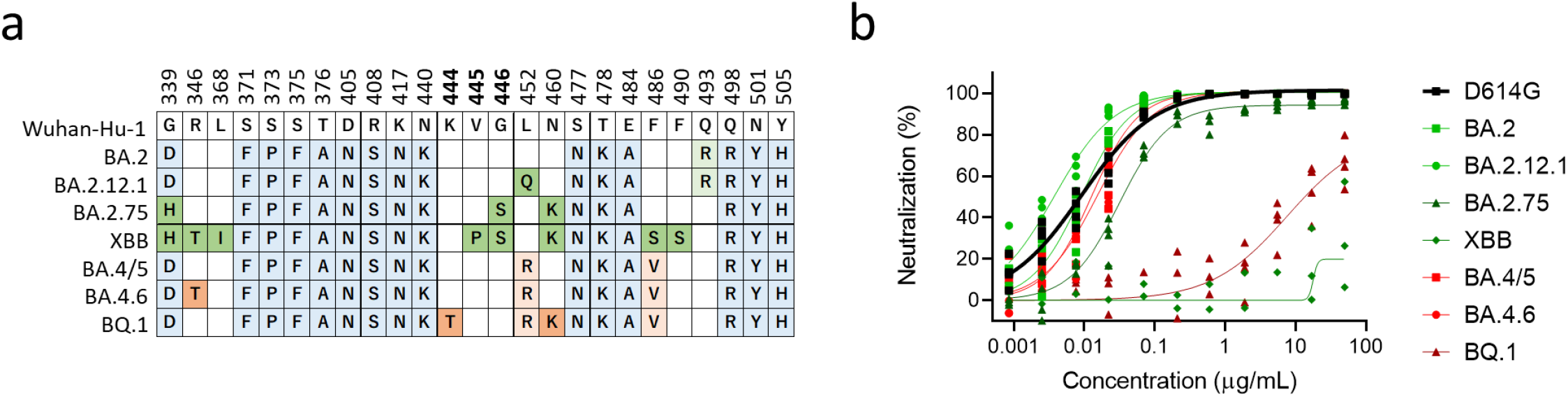
Bebtelovimab neutralization efficacy against the Omicron subvariants. (**a**) Location of mutations in the receptor binding domain proteins of the Omicron subvariants; BA.2 and its derivatives, BA.2.12.1, BA.2.75, XBB, and BA.4/5 (which are identical at the amino acid level) and its derivatives, BA.4.6, and BQ.1.1. Conserved mutations among omicron subvariants are highlighted in blue. Unique mutations in BA.2 derivatives and BA.4/5 derivatives are highlighted in green and red, respectively. Top number is the residue according to the spike protein of SARS-CoV-2 Wuhan-Hu-1. (**b**) Neutralization efficacy of bebtelovimab in 293T/ACE2 cells. n = 4 technical replicates.

## Discussion

After emergence of SAAES-CoV-2, therapeutic monoclonal antibodies (mAbs) were rapidly developed and clinically used to prevent both infection and disease progression(14). In the COVID-19, virus neutralizing mAbs has become the popular anti-viral strategy. However, mutational escape of virus from monoclonal antibodies were observed from the early period on the clinical use. Bamlanivimab was the first mAb granted Emergency Use Authorization (EUA) to treat COVID-19 in November 2020 by the U.S. Food and Drug Administration (FDA) but the spread of E484K-carrying variants led to its withdrawal in April 2021(15). To prevent such viral evasion, some mAbs were used in cocktail form and others were designed to target highly conserved regions(11, 16, 17). However, considerable antibody escape was observed in the Omicron and continuously emerging subvariants keep threatening the efficacy of therapeutic mAbs. Thus, it is crucial to comprehensively understand the antibody escape to effectively manage to effectively manage future variants.

DMS is the powerful tool to understand the functional alteration due to each amino acid substitution and applied to predict antibody escape. One approach to evaluate the antibody escape is the assay for antibody binding using yeast-surface display of the RBD domain and separately assess the ACE2-binding that reflect the infectivity(18, 19). Instead, we originally developed the assay system that reproduce the viral infection. The full-length spike protein was expressed on human Expi293F cells and incubated with ACE2-harboring pseudotyped lentivirus. This assay mimics well the infection that is the fusion of cell and viral membranes mediated by protein-protein interactions between virus spike and ACE2, albeit in an inverted orientation. When examined with antibodies, this assay can directly the viral escape that is the results of antibody-binding and infectivity.

Crystal structure analysis provides the information of antibody epitope. However, not all epitope sites and not all amino acid substitutions are responsible for antibody escape (Figure 1e). DMS can comprehensively identify the antibody escape and this detailed characterization leads to the effective management of future SARS-CoV-2 variants.

## Methods

### Cell culture

HEK 293T cells (Clontech) and its derivative, 293T/ACE2 cells were cultured at 37 °C with 5% CO_2_ in Dulbecco’s modified Eagle’s medium (DMEM, WAKO) containing 10% fetal bovine serum (Gibco) and penicillin/streptomycin (100 U/ml, Invitrogen). All cell lines routinely tested negative for mycoplasma contamination.

### Protein synthesis and purification

The sequence of bebtelovimab was obtained from KEGG DRUG Database (https://www.genome.jp/kegg/drug/) and formulated in the form of human IgG1/kappa by using synthetic DNA coding for the variable regions of heavy and light chains taken from the publicly available amino acid sequences, and recombinantly produced in Expi293F cell expression system (Thermo Fisher Scientific) according to the manufacturer’s protocol. Fc-fusion proteins were purified from conditioned media using the rProtein A Sepharose Fast Flow (Cytiva). Fractions containing target proteins were pooled and dialyzed against phosphate buffered saline (PBS).

### Deep mutational scanning for escape from monoclonal antibodies

Monoclonal antibody selection experiments were performed in biological duplicate using a deep mutational scanning approach with previously described mutant RBD libraries(1). The library was focused on the BA.2 strain spike RBD region from F329 to C538. Pooled oligos with degenerate NNK codons were synthesized by Integrated DNA Technologies, Inc. Synthesized oligos were extended by overlap PCR and cloned into pcDNA4TO HMM38-HA-full length spike plasmids. Transient transfection conditions were used that typically provide no more than a single coding variant per cell(20). Expi293F cells at 2 × 106 cells per ml were transfected with a mixture of 1 ng of library plasmid with 1 μg of pMSCV empty plasmid per ml using ExpiFectamine (Thermo Fisher Scientific). Twenty-four hours after transfection, cells were incubated with human ACE2 (hACE2)-harboring green fluorescent protein (GFP) reporter viruses, which were generated by transfecting pcDNA4TO hACE2, psPAX2 (addgene #12260), and pLenti GFP into LentiX-293T cells with Lipofectamine 3000 (Invitrogen). The viruses carry hACE2 instead of glycoprotein can infect the cells expressing the spike. To analyze the escape mutations from antibodies, cells were pre-incubated with antibodies for 1 hour. Next, these cells were treated with hACE2-harboring virus for 1 hour and further incubated with fresh medium for 24 hours. Cells were harvested and washed twice with PBS containing 10% bovine serum albumin (BSA) and then co-stained for 20 minutes with anti-hemagglutinin (HA) Alexa Fluor 647 (clone TANA2, MBL). Cells were again washed twice before sorting on a MA900 cell sorter (Sony). Dead cells, doublets, and debris were excluded by first gating on the main population by forward and side scatter. From the HA positive (Alexa Fluor 647 positive) population, GFP-positive and -negative cells were collected. The total numbers of collected cells were about 2 million cells for each group. Total RNA was extracted from collected cells using TRIzol (Life Technologies) and Direct-zol RNA MiniPrep (Zymo Research Corporation) according to the manufacturer’s protocol. First-strand complementary DNA (cDNA) was synthesized with PrimeScript II Reverse Transcriptase (Takara) primed with a gene-specific oligonucleotide. The RBD library was divided into 3 sections with 210bp fragment each and pooled before transfection to analyze the same experimental condition. After cDNA synthesis, each library was amplified with specific primers. Following a second round of PCR, primers added adapters for annealing to the Illumina flow cell, together with barcodes for each sample identification. The PCR products were sequenced on an Illumina NovaSeq 6000 using a 2 × 150 nucleotide paired-end protocol in the Department of Infection Metagenomics, Research Institute for Microbial Diseases, Osaka University. Data were analyzed comparing the read counts with each group normalized relative to the wild type sequence read-count. Log10 enrichment ratios for all the individual mutations were calculated and normalized by subtracting the log10 enrichment ratio for the wild type sequence across the same PCR-amplified fragment.

### Pseudotyped virus neutralization assay

Pseudotyped reporter virus assays were conducted as previously described(1, 21). Using a plasmid encoding the SARS-CoV-2 spike protein (addgene #145032) as a template, D614G mutant and Omicron subvariants were cloned into pcDNA4TO (Invitrogen) in the context of the ΔC19 (19 amino acids deleted from the C terminus) (22). Spike protein-expressing pseudoviruses with a luciferase reporter gene was prepared by transfecting plasmids (pcDNA4TO Spike-ΔC19, psPAX2 (addgene #12260), and pLenti firefly) into LentiX-293T cells with Lipofectamine 3000 (Invitrogen). After 48 hours, supernatants were harvested, filtered with a 0.45 μm low protein-binding filter (SFCA), and frozen at −80 °C. The 293T/ACE2 cells were seeded at 10,000 cells per well in 96-well plates. Pseudoviruses and three-fold dilution series of antibodies were incubated for 1 hour, then these mixtures were added to 293T/ACE2 cells. After 1 hour incubation, the medium was changed. At 48 hours post infection, cellular expression of the luciferase reporter, indicating viral infection, was determined using ONE-Glo Luciferase Assay System (Promega). Luminescence was read on Infinite F200 pro system (Tecan). This assay was performed in four replicates and the 50%-neutralizing concentration was calculated using Prism version 9 (GraphPad Software).

### Statistical analysis

Pseudoviruses neutralization measurements were done in technically four replicates and relative luciferase units were converted to percent neutralization and plotted with a non-linear regression model to determine 50% neutralizing concentration using GraphPad Prism software (version 9.0.0).

## Acknowledgements

We would like to thank Yuki Ozaki and Daisuke Motooka (Department of Infection Metagenomics, Research Institute for Microbial Diseases, Osaka University.) for sequencing of DMS samples.

## Author contribution

A.H. and S.M. designed the research; Y.H. performed pseudovirus neutralization assay; M.C.A. and Y.H. performed deep mutational analysis; Y.K. performed and analyzed next-generation sequencing; M.C.A. purified and prepared the proteins; A.H. wrote the manuscript; all authors discussed the results and commented on the manuscript.

## Data and materials availability

Deep mutational scan data has been deposited at SpikeDB (https://sysimm.ifrec.osaka-u.ac.jp/sarscov2_dms/).

